# Reconstructing Biological Molecules with Help from Video Gamers

**DOI:** 10.1101/2024.06.19.599674

**Authors:** Andreas C. Petrides, Robbie P. Joosten, Foldit Players, Firas Khatib, Scott Horowitz

## Abstract

Foldit is a citizen science video game in which players tackle a variety of complex biochemistry puzzles. Here, we describe a new series of puzzles in which Foldit players improve the accuracy of the public repository of experimental protein structure models, the Protein Data Bank (PDB). Analyzing the results of these puzzles showed that the Foldit players were able to considerably improve the deposited structures and thus, in most cases, improved the output of the automated PDB-REDO refinement pipeline. These improved structures are now being hosted at PDB-REDO. These efforts highlight the continued need for the engagement of the lay population in science.

## Introduction

Our shared knowledge of experimental biomolecular structures is stored in the Protein Data Bank (PDB) (*1*). The accuracy of the PDB is important for multiple reasons. For example, the recent successes in deep learning approaches to predicting protein structure (*2*) rely on an accurate PDB for continued development. Similarly, the PDB serves as a resource for biologists and biochemists to develop hypotheses to test in their everyday experiments. Mistakes in structure models can cause scientists to base their hypotheses on incorrect data, which causes delays in scientific progress.

The PDB contains many solvable errors (*3*). The type of data used in crystallography and cryo-EM, maps of electron density or electron potential, respectively, have limitations as they are the result of an indirect experiment with experimental error and limited resolution. By showing where the electrons are, scientists can fit atoms to these maps to discover the atomic structure of the macromolecule in the experiment. This requires a combined approach of visual fitting of these maps as well as computational tools to aid in complying with known rules of chemistry and physics in how molecules are put together. Limitations include: low resolution of the data, parts of the maps where artifacts or missing data can cause blurriness in the maps, assumptions in computational models used to process the primary experimental data, among other causes of low data quality. As a result, the scientists interpreting these data can make mistakes. High attention to detail and the use of verification tools can help prevent some mistakes (*4*), but the sheer quantity of data makes it likely that human error will still persist into published and deposited structure models. Furthermore, the PDB does not require peer review for deposited models, and as a result, many entries that contain errors ranging from inconsequential to egregious exist within the PDB.

For this reason, the PDB-REDO project was begun in 2006 (*5*). The mission of PDB-REDO is to perform automated re-refinement and rebuilding of the crystallographic structure models in the PDB to improve the accuracy of the PDB and remove model errors in the process. This venture has been highly successful, with the PDB-REDO databank now containing over 170k entries, many of which have an improved fit to the experimental data and more probable structural features (*6*).

However, is there still a need for improvement of structure models beyond the automated re-refinement by PDB-REDO? Yes, the procedure in PDB-REDO looks at many structural aspects (*7*) but is limited to model issues that can be handled robustly with a very low risk of making the model worse. Indeed, a search of the PDB for its lowest quality structure models shows that the improvements by PDB-REDO in these cases can be modest at best. They still appear to be relatively low quality after PDB-REDO refinement. As a result, new approaches are needed.

One possible approach is to enlist the aid of humans to improve the PDB via the players of the biochemistry video game Foldit. Foldit is a citizen science game in which players work on a variety of complex biochemistry puzzles, collaborating and competing to create the best possible structure model (*8*). Model quality within the game is judged by the Rosetta force field (*9*), combined with other elements, such as the fit to an underlying map from experimental data (*10*). Previous competitions have shown that Foldit players can solve both crystal structures and cryo-EM structures with higher accuracy than scientists or computational algorithms (*10, 11*). Therefore, it became an obvious question to ask whether Foldit players could also improve the structures already within the PDB?

## Results

To test this possibility, we created a new series of Foldit puzzles, termed *Reconstruction puzzles*. In these puzzles, Foldit players were given protein structure models and the underlying density maps and tasked with improving the structures. In this report, we discuss the results after the first 58 of these puzzles.

After the puzzles, the players’ solutions were clustered and then analyzed by PDB-REDO for comparative analysis to see if using the Foldit structure models as a starting point is an improvement over using the structure models from the PDB itself. In 39 of the 58 cases, the overall structure quality was improved by using the Foldit structure. In general, the Foldit player structures had considerably improved chemical and physical properties in the vast majority of puzzles by multiple metrics, and in many cases also improved the fit to the experimental data (Fig 1 and Supplemental Table 1). As seen by Molprobity ClashScore (*12*) (Fig 1C), the atomic clashes of the structure models were especially improved. Two examples of before and after the Foldit use can be seen in Fig 2. In some cases, such as in Fig 2A, changes to the structure can be observed at many points. In many of the cases though, such as in Fig 2B, the changes are localized, and often in regions of poor electron density such as loops. However, these regions can be important biologically. For example, in the case of 2fxt, yeast mitochondrial import inner membrane translocase subunit TIM44, we observe that Gly388 has changed position by 3.2 Å due to a so-called register shift. This glycine is fully conserved over 1522 sequences in HSSP (*13*) and in the more distant human ortholog (Uniprot AC: O43615) this glycine is conserved as well. Moreover, AlphaMissense (*14*) suggests that mutations to this residue are very likely to be pathogenic (minimal score 0.922) indicating that this a key residue in the protein.

**Figure 1:**
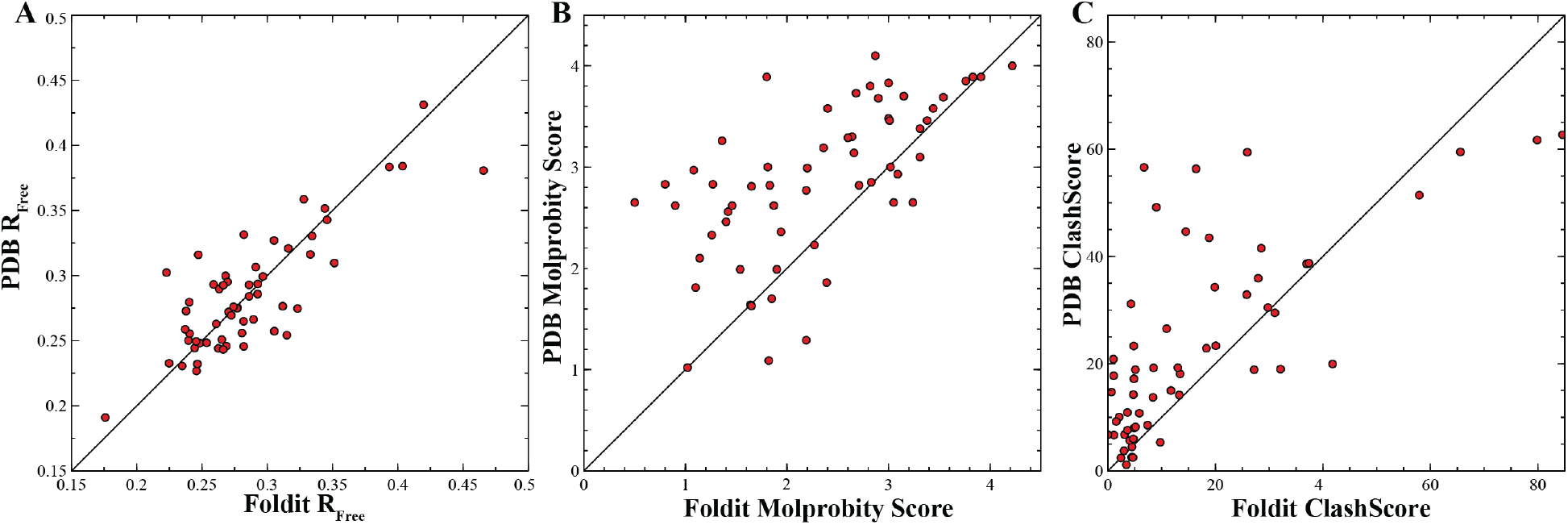
Comparison of selected statistics for PDB-REDO output using original PDB entry (y-axis) or Foldit-refined entry (x-axis) as input. Each point is a separate PDB entry. A. R_Free_, B. Molprobity Score, and C. Molprobity ClashScore for each entry. As lower scores are better, above the diagonal line are cases in which the Foldit model improved over the original PDB entry as the starting point for PDB-REDO.

**Figure 2.**
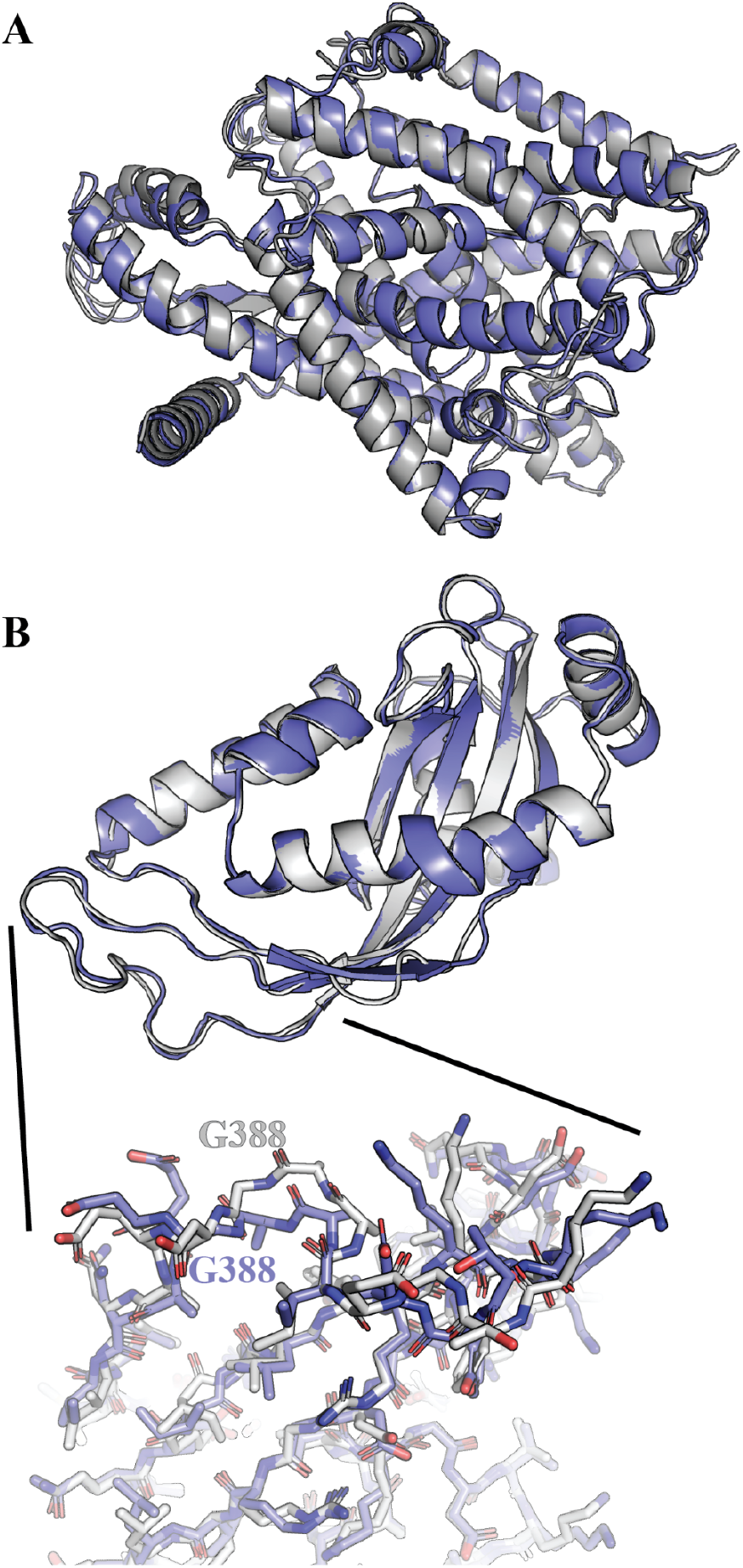
Structure of 2yxq (A) and 2fxt (B). Foldit structure used as starting point for PDB-REDO in blue, PDB-REDO using original PDB as starting structure in gray.

With the success of the *Reconstruction puzzle* series, we have created a new pipeline in which data from Foldit puzzles can be automatically analyzed and evaluated by PDB-REDO for inclusion in the PDB-REDO database. In these cases, when the Foldit player has agreed to have their Foldit name attached to the entry, the PDB-REDO webpage will give credit to the Foldit player whose structure model is now available for public download.

How did the Foldit players improve on these structures? To get an idea, we asked several Foldit players whose structure models were chosen as the best overall structure. The approaches taken by different players was quite variable. In most cases, the strategy involved a combination of hand-fitting as well as automated tools and scripts. However, in certain cases, the top solutions used only scripts and automated tools, suggesting that considerable improvement in automated fitting of structural maps can be accomplished (See Supplemental Information).

## Conclusions

With the success of the Foldit players in improving experimental structure models, the *Reconstruction Puzzle* series within Foldit continues, and will continue to provide improvements to known structures. At present, these puzzles are only for crystallographic structure models, but with the increase in number of cryo-EM structure models, the same problems persist there as well, and can also use the intervention of Foldit players.

With the focus on new Artificial Intelligence tools and how they can improve fields including structural biology, cases like this are here to remind us that the greatest untapped potential for science is that of humanity.

## Methods

*Reconstruction puzzles* were prepared by performing 5 rounds of refinement in Phenix (*15*) using the original PDB entry with all ligands and waters removed and including simulated annealing. This new model was given to the players with a Feature Enhanced Map (*16*) for reconstruction.

### A set of post processing steps occurred following each puzzle’s completion

Using retrieved player-generated solution files, a Python script orchestrated the clustering of PDB files based on their structural similarity, employing the Cα Root Mean Square Deviation (CA-RMSD) metric. Iteratively, the script varies the threshold for CA-RMSD clustering, starting from 1 Å and decreasing until the total number distinct clusters is greater than 100. This iterative clustering process is crucial since player-generated solutions contain a diverse set of structural configurations. This set of steps filtered the structures down into a concise set of models.

Following clustering, the solutions were clustered and ranked based on their Foldit score, with the top 100 clustered solutions moving to the next step. The Python script then submitted the top 100 clustered solutions by score to the API of the PDB-REDO web service (*17*), facilitating the submission of these clustered solutions for further refinement and analysis.

The PDB-REDO procedure used here entailed optimizing the weight between the experimental data (as retrieved from the PDB) and restraints for covalent geometry and atomic B-factors to maximize the fit with the experimental data whilst maintaining normal geometry. Combinations of different types of additional restraints were used while refining the models with REFMAC (*18*): homology-based hydrogen bond restraints (*6*) were applied with 8 solution sets, general hydrogen bond restraints with 18 sets, local non-crystallographic symmetry restraints with 26 sets and jelly-body restraints with 19 sets (*19*). All the model rebuilding steps in PDB-REDO were excluded to stay close to the original solution.

The Python script awaited confirmation of completion for each solution from the PDB-REDO API. Upon completion, the finalized protein structures and associated metrics were retrieved. Deposited structures were written into new PDB files. Corresponding metrics were organized into comprehensive data dictionaries and graphs, both of which facilitate further analysis.

## Acknowledgements

We would like to thank J. Flatten for his assistance maintaining the Foldit platform and the RHPC facility of the Netherlands Cancer Institute for maintaining PDB-REDO’s compute infrastructure. Funding was provided by NSF 2051305 to SH, NSF 2051282 to FK, Horizon 2020 research and innovation program grant No. 871037 (iNEXT-Discovery) to RPJ and by an institutional grant of the Dutch Cancer Society and of the Dutch Ministry of Health, Welfare and Sport.

The authors declare that they have no conflicts of interest.

## Supplemental Information-Foldit Player ED strategies

### NinjaGreg

1 I begin by setting the wiggle power to Low. This gives the segments the most flexibility to adjust their shape.
2 Since we can assume that the initial fold of the protein is pretty close to what it should be, I don’t apply any hand folding. The first thing I do is get the protein as relaxed as possible.
3 I “shake” until it has made at least two passes over the protein. This receives a lot of the stresses that occur as part of importing the protein into Foldit.
4 I “wiggle” until the score stops changing. This also releases much of the stresses from the import.
5 I try another shake. If it gets points, I repeat steps 3 and 4 until no more score gain is achieved.
5 I then run the script “cut and wiggle”, which cuts the protein every N segments, then wiggles all segments. This will allow the segments to move more freely, and will release more of the stresses in the segments.
6 I then run the script “micro idealize”, which applies a cut to one end of each segment, then “idealizes” that segment, then removes the cut and wiggles that and a few surrounding segments.
7 I then run the recipe “QuakeR”, which uses bands to apply compression to selected segments, looking to “squeeze out” any avoidable voids inside the protein. It’s sort of like squeezing a Brillo pad to reduce its volume. At this point the protein is pretty relaxed, so we can start seeking to improve the configuration. If disulfide bonds need to be connected, this is the point to do it. I use hand folding with bands, it’s pretty fast.
8 I normally then run the script “fracture”, selecting the “early” option, using the first option presented. I run this overnight, allowing the script to hopefully examine the entire protein at least once. This option of fracture works on first three segments, then one more each pass up to six segments, rebuilding the selected segments multiple times, and using the highest scoring rebuild if it improves the overall protein score (otherwise it reverts back to the positions at the start of that set of segments.)
9 I then run the script “AFK3”, which reduces the clashing importance, wiggles until the score stops declining, shakes the protein to relax the side chains into more favorable positions, raises the clashing importance back to full and wiggles until the score stops improving. It may repeat this process with a smaller reduction in the clashing importance to see if it can improve the score still more. It repeats this process until it fails to improve the score a given number of times (or until I see it not making progress, at which point I cancel it).
10 I may then run other scripts to continue to wring points out of the protein, which means that the energy of the protein is being reduced. Typical programs I run at a this point are “JET”, which does wiggling in sets of segments of various lengths to look for more points (less energy); “fuze using cuts”, which places a cut point at selected places, does a “fuze”, removes the cut, and “fuzes” again; “TvdL enhanced DRW”, which is similar on concept to fracture, but investigates rebuilds based on a list of different criteria; “nbl_hinge”, which uses a variety of subscripts to massage the protein; and sometimes “Random idealize”, which randomly selects groups of segments and idealizes them, then wiggles the result to see if the score is improved.

I run the various scripts until it appears that as much improvement as possible has been achieved. I then set the Wiggle Poser to High, and repeat the above sequence.

The End Game

Once the above sequence has squeezed the protein until little more progress is made, then I resort to running first “Banded Worm Pairs Infinite” with wiggle power set to Low, then “AFK3” with wiggle power set to High. I repeat these two until they stop yielding points, then try various of the above scripts to seek further improvement, doing this until time runs out on the puzzle (usually one week).

It is particularly important to do as much score improvement as possible with the wiggle power on Low in the beginning as this is when the protein is most relaxed and susceptible to movements. Once the switch to High wiggle power has been made, the protein gets “stiffer”, harder to reshape.

### MicElephant

I currently have a typical sequence that I use for the smaller ED puzzles. First steps all with wiggle power auto.

1) Wiggle sidechains, shake sidechains, wiggle all. Repeat this sequence until no more gains.

2) Tvdl Walking Rebuild, length 1 to 5 for smaller puzzles, about 12 hours for the real big ones.

3) Tvdl remix, length 3 to 6, until no more gains for ∼ 1 hour. Sometimes I use Jolter 2.0 instead.

4) Microidealize 4.1.1

5) Cut&wiggle everything, length 3

6) Tvdl DRW compressor 2.0, length 2 to 6, until no more gains for about an hour.

7) Sidechain flipper, using extensive mode

8) nbl_hinge 3.4 with default settings

9) Total LWS 2.0

Now wiggle power medium

10) Wiggle all

11) AFK 3.1

Now wiggle power high

12) Wiggle all

13) AFK 3.1

Now wiggle power low

14) BandedWorm Pairs IntFilt 1.4.9 until 5 to 10 points are gained

Back to wiggle power high.

15) From here on: no fixed sequence. Repeat one of the recipes mentioned above.

16) Repeat step 14 (low) and 15 (high)

Sometimes if I see bad parts of the protein which are not fixed by the recipes I do some hand folding between the steps, e.g. to get sheets more parallel, or if a sidechain is not in the density cloud at all.

### Hillbillie

First of all, my approach to the mentioned puzzle is the same that I use for all puzzles with all work done by self programmed lua scripts. Since I have no background in protein construction or manual improvements to these structures, I simply started to use scripting to do all of my work.

The main indicator in this working process is the total score after different types of interactions with the structure and using most of the capabilities of the given scripting commands. I’m still trying to improve my results by adjusting my scripts and letting them do their work automatically as long and as fast as possible. The choice which kind of interaction to be done is mainly based on random selection.

So you can say my results are based on lucky guessing and finding best possible ways of improvement.

This might not sound very groundbreaking, but that’s my way of working through the puzzles and trying to create powerful algorithms. I hope I could give you a brief look at my work.

### Bruno Kestemont

The good news is that for these revisited puzzles, I don’t hand fold nor do I show the ED map: everything is done with recipes. This suggests that everything could be automated for a mass treatment.

I will start by explaining my overall strategy, then I document what I did for the specific puzzles.

#### Overall strategy

I always set wiggle power to low. When the deadline is <3 days, I turn to high wiggle power.

I usually start 3-5 different tracks on the same puzzle. It depends on the available memory. I start with “auto structure”.

On different tracks, I run shake-wiggle sidechains. One track with wiggle sidechains-shake-wiggle sidechains.

Then I start with a different recipe on each track, overnight.

I always use the following 2 recipes within the period of the first 4 days on low WP: - https://fold.it/recipes/102471 jolter 2.4 (in short: j), days 1 to 3, 8-24 hours running. I can stop it after 12 hours, run auto-structure and start it again, until there is no more gain. When hand folding on previous ED puzzles from scratch, I observed that a secondary structure that was already roughly hand positioned in a cloud would then be finely positioned with full (helix/sheet) rebuild and in particular this recipe. Thus, this is a perfect recipe for refining ED puzzle solutions. Also, the loops find a reasonable position in the cloud with this (if there is no time left and the helices and sheets are already well positioned, I could focus the recipe on loops only). The recipe rebuilds all identified SS and loops in turn infinitely.

-https://fold.it/recipes/100947 Cut and Wiggle Random v1.1 (in short: cwr), day 1 to 4 (preferably after other recipes because it tends to stick the protein), 8-24 hours running, until it gains less than several points per loop.

When hand folding on previous ED puzzles from scratch, I observed that a rough hand positioning of a cut piece in the cloud “automatically” perfectly fitted the section in the cloud with wiggle all. Thus cut a section, put in the cloud, wiggle all this section. I positioned the helices and sheets piece after piece like this. When I was finished, I closed the cuts before to wiggle again.

Thus, Cut and Wiggle Random is an excellent recipe for fine tuning an already solved ED puzzle.

Note that this gives me an idea for a new recipe combining Jolter and Cut & wiggle (cutting before and after a SS before to wiggle, then uncut). Thus cut & wiggle SS (not random). To code asap.

If there is time left, I then run the following in order to rebuild low density score segments. -https://fold.it/recipes/105055 Fracture v2.1 w/ Remix (in short: fr or frw). Day 2-4, 8-16 hours. The options are set as follows: “Early game - Fast & Loose”, “I’ll have banders with that”, “Do remix instead of rebuild …” (if there is enough time left), and certainly “Target density subscore”.

If there is time left, I could run a recipe including random idealize function, like https://fold.it/recipes/49751 Quaking Rebuild V2 1.0 RI (in short qrri)

The other recipes I could run as first recipe before j and cwr are: https://fold.it/recipes/102831 AFK3.5.1 (BounceWiggle). In short afk (with no shake) or afks (with shake) https://fold.it/recipes/108480 Quickfix 3.6.3 + faster cleanup (in short: qf)

The second recipe (day 2-4) is always J, third recipe is always cwr. Fourth recipe is fr if there is time left.

Then if there is time left, I run a local wiggle recipe in order to refine before to switch to high wiggle power: -https://fold.it/recipes/103159 JET 4.2.7

Other recipe I could use on day 3-4 before jet if there is time: https://fold.it/recipes/101746 Ebola 4.3 (option velocity 5) (it rebuilds the worst long portions)

Then, switch to high wiggle power and run the following recipes: -https://fold.it/recipes/103121 nbl_hinge_v4.2.2 (in short nblh) -https://fold.it/recipes/108343 Banded Worm Pairs Inf Filt 3.5.7 (in short bwpi)

#### Specific puzzles

https://fold.it/puzzles/2013511 (2224)

Auto structure, shake, wiggle sidechains

On low wp: afks + fr + j + cwr

On high wp: high nblh + auto bwpi then several recipes in high wp in parallel with auto bwpi (in parallel means that several tracks run the same solution in parallel, saving and loading regularly from local share in order to start over with the best one on all tracks).

https://fold.it/puzzles/2013617 (2300)

It was a small puzzle, I could use my full range of favorite ED recipes.

Auto structure, shake, wiggle sidechains

On low wp: j + cwr + j + qf + qrri + fr + ebola + cwr + jet

On high wp: high nblh + auto bwpi then several recipes in high wp in parallel with auto bwpi

https://fold.it/puzzles/2013706 (2359)

Auto structure, change segments to helix and idealize this portion, shake, wiggle sidechains

On low wp: (probably cwr +j+ fr) + ebola

On high wp: high nblh + auto bwpi then several recipes in high wp in parallel with auto bwpi

https://fold.it/puzzles/2013714 (2368)

Auto structure, shake, wiggle sidechains (or reverse, I don’t remember)

On low wp: (probably j+ cwr + fr) + qrri + ebola + cwr + jet

On high wp: high nblh + auto bwpi then several recipes in high wp in parallel with auto bwpi

### blazegeek

I’ve been struggling to find a way to describe my approach as I don’t really have one. The only consistent pattern I notice is that I try several different approaches simultaneously until there is an obvious score leader. I then try several approaches using that solution until, again, there is a clear leader.

I will point out that I don’t have a background in molecular biology or related fields (I actually studied television production). In fact, I still have a hard time remembering the 20 amino acids without forgetting one or two. I suspect that my lack of knowledge and understanding enables me to try things that most others wouldn’t consider trying, resulting in a sort of “beginner’s luck” on occasion.

Being neurologically atypical is certainly a factor in my approach to problem-solving of any kind, but especially within the highly stimulating, interactive environment of Foldit. Unfortunately, it would be impossible to describe what that process entails. I tend to look at a protein aesthetically; some shapes are much more pleasing than others, more beautiful, more elegant. It’s useful to be able to visualize this way, as that elegance seems to exist throughout Nature and at all scales. It may not be very scientific, but one doesn’t need to understand something to find it beautiful. That, too, is rather elegant in itself.

I don’t know how helpful this is for your paper, but I appreciate the opportunity to share my approach. In a nutshell, I tend to approach these puzzles quite abstractly, which occasionally pays off. I’m happy that something useful came of it!

### spmm

My general approach: try to identify any large hydrophobics, helices and the ‘ends’ in the density cloud, attempt to understand where the ‘folded mass’ is located and assume I will need to start again once a picture emerges.

### gmn

No problem sharing my approach. It hasn’t changed much if I’m using the trim tool. Approach depends on the size of the protein.

### Very large ED proteins (multiple subunits)

Start with CI 1.0/Low wiggle

Shake/wiggle

Fix any bad scoring areas with remix (usually the red segments) or rebuild if remix doesn’t work

Freeze entire protein

Unfreeze only the subunit I want to work on and highlight it for trim tool

Rebuild trimmed subunit with scripts

Untrim/shake/wiggle

Unfreeze entire protein

Shake/Wiggle (compare to untrimmed s/w score and take higher score for next subunit)

Freeze entire protein/unfreeze subunit I rebuilt/highlight for trim tool/run GAB script on trimmed subunit

Untrim/shake/wiggle

Unfreeze entire protein/shake/wiggle (compare to s/w untrimmed score and take higher score for next subunit)

Repeat process for each subsequent subunit I want to work on

Run end game scripts on entire protein, which includes switching to high wiggle at some point, once I have completed working on all submits

### Smaller ED proteins (few or no subunits)

Start at 0.5 CI/Low wiggle

Shake/wiggle

Fix any bad scoring areas with remix (usually the red segments) or rebuild if remix doesn’t work

Freeze entire protein

Unfreeze only the section I want to work on and highlight it for trim tool Rebuild trimmed area with scripts

Untrim/shake/wiggle

Unfreeze entire protein

Shake/Wiggle (compare to untrimmed s/w score and take higher score for next section) Repeat process for each section I need to work on (Note: I do not usually GAB at CI 0.5.) Repeat the entire process for CI 1.0 (see above)

If the ED puzzle is sufficiently small, I will start at CI 0.2 and run the CI 0.5 process using CI 0.2 (no GAB). I will then follow up with the CI 0.5 process and add GAB for each section (which then leads to the CI 1.0 process).

### LociOiling

I use the same general approach on all electron density puzzles. My approach involves running different sequences of recipes on two or more parallel tracks.

Many of the recipes are private (or group) recipes, but all are based on public recipes.

The initial steps are always done on low wiggle power. I shift to high wiggle power later, roughly halfway through the week that the puzzle runs.

The first step is to “wiggle out” the starting pose. I first shake (S), then wiggle sidechains (E), then wiggle all (W). I usually let the wiggle steps go until the counter spins. I repeat the SEW action two or three times.

At this point, I “nudge” the protein by pulling on a segment to make the score drop. I wiggle again to see if the score recovers. I repeat SEW if it reaches a higher score. I may also use a band plus wiggle to drop the score, then delete the band to see what happens.

After the score reaches a plateau, I run a Fuze recipe, Fuzes 3.0.3, usually using the default settings.

After Fuzes, I have the starting point that I’ll use on the rest of the puzzle. I usually take a look at the PDB at this point, to see if I can match the protein.

Taking the starting point solution, I start two or three clients. Each client runs in its own directory, minimizing any interaction.

Still on low power, one client runs TvdL Enhanced DRW 3.1.1 (EDRW), which uses the rebuild function. I generally increase the number of rebuilds per section, from the default of 15. For a small puzzle, I would go with 100 rebuilds per section, but I set a lower number for monster puzzles. The number of cycles still defaults for 40, but I always reduce this to eight or less based on the size of the puzzle.

A second client runs a private remix recipe based on TvdL DRemixW 3.1.2.

I may also try a third client, using a slight modification of the public BandFuze recipe. This recipe keeps running until cancelled. Sometimes I may remix or rebuild the results.

After the remix completes, I switch to rebuild, and rebuild switches to remix. With the trend toward very large puzzles, I may reduce the number of cycles at this stage.

Once remixing/rebuilding is complete, I switch to high wiggle power. The exact timing of course depends on the size of the puzzle. Usually the rebuild/remix and remix/rebuild solutions are still in the running at this point. The BandFuze solution may also still be viable.

The first recipe on high power is a private “GAB” banding recipe. I adjust the number of GAB generations based on the puzzle size.

After GAB, I run a Cut & Wiggle recipe followed by a Microidealize recipe. My versions are more persistent than most of the public versions.

After Cut & Wiggle and Microidealize, I switch to an LWS recipe, still on high wiggle power. I use a private recipe that’s based on Band Worm Pairs to start.

When my private recipe begins to plateau, I usually switch to auto wiggle power, and run Banded Worm Pairs Inf Filt 3.4.10 (BWP IF) by Bruno Kestemont.

At this late stage, I also assess the competition. I may continue with an LWS recipe if I’m in the lead. Sometimes, corrective action may be helpful, such as additional remixing on rebuilding on high wiggle power. An Acid Tweaker recipe can sometimes help. There are also a number of other LWS recipes on the bench, and sometimes they can find a point or two.

I’’m working on improving EDRW and other key recipes to make them “chain aware”, since many of the recent ED puzzles have two or more chains. I’m guessing that remixing across chain ends is never a good idea, but it’s not clear how much time it actually wastes. A similar argument applies to rebuild and other actions.

### A quick followup to the previous email

I looked at the saved solutions for the six puzzles listed, and didn’t find anything too different than what I previously stated. On one of the puzzles, an Acid Tweeker near the end found a large gain. Otherwise, the process seemed to be much as described.

I didn’t mention that after running Band Worm Pairs on auto for a while, I may switch back to high wiggle power if time permits. Going back to high often finds a few points.

I also didn’t mention the trim tool. I’ve tried it on several puzzles with mixed results. On puzzle 2439, I tried remixing and rebuilding adjacent pairs of chains. This still took forever due to the large number of chains. The score was lower after finishing each pair. The winning solution was based on tackling the entire protein.

On other puzzles, working on trimmed sections separately has produced gains instead of losses. I think the “everything all at once” strategy was still the winner in most of these cases.

The trim tool itself seems to be working well, so possibly a better strategy is needed. Currently, I’ve been shaking, shaking, wiggling and fuzing after un-trimming each section. Maybe saving these steps until all sections have been processed would work better.

### toshiue

I found my notes on that and as I had expected, there wasn’t anything unusual (for me) in the approach to that protein. That said, I don’t have visibility on the approaches taken by others, only those that I’ve done on EDs in the past. You have a greater sampling to compare such to and my notes are four pages, if you’d like a copy.

I/we bump up against and sometimes exceed the theoretical limits of a protein. I think 2300 was just one of those instances where more things went right than usual. I don’t have a rigid approach to folding. I primarily do evo only with enough solo to get a taste for the protein.

To me, each protein is an individual and I approach them as such. Group dynamics plays a greater role than one might imagine. When most of our group veterans are present and we’re slightly giddy, we are truly at our best as a group. I returned to folding when I noticed the EDR series had continued (2300 being one of the EDR series). Such are very easy, for me, to work with and I thoroughly enjoy them. Thus, the hours get put in on such. On that note, putting in the hours is a necessity. Hardware matters more than most seem to believe. Iterations matter.

I run clients on all three major OS, Windows, Mac, Linux. Recipes respond differently to each. Often when we push a protein hard on one of our favorite (demanding) recipes, we get swarms of crashes. Switching over to another OS very often cures this. One last item, on the EDR series. They’re too anonymous. Descriptors such as the one on EDR88 don’t entice us into all nighters. Adding real world descriptors (ie… It’s a delivery system for a new pediatric cancer drug at Stanford Medical), and we’re pulling all nighters.

Sorry I didn’t have a magic algorithm to offer, but these proteins really are individuals. We have to get to know them before we know what we can do with them. Also, I think too many are overly cautious. There’s a cultural norm as to approaches with the folding regimen that everyone seems to strive for. I routinely break with those norms.

### Galaxie

I start by looking at the structure to see how it fits in the ED. If DNA is present, I try to get the bonds in line for the correct pairs.

Starting with low wiggle and progressing to auto wiggle I use a series of rebuilders and banders. Cut and wiggle, more banders and microIdealize are used in high wiggle. End game usually involves banded worm pairs. Sometimes rebuilders and acid tweaker are helpful to further refine the structure.

### AlphaFold2

#### Just a few simple tips on Electron Density Reconstruction Puzzles

Before I start to use any tools, I will rotate the Protein 360 degrees to see if there are any areas that stand out from the rest. Patterns, I identify these by looking for amino acids that are not fully extended or hydrophilic or hydrophobic and any segment of the backbone that may have a better potential in a new configuration.

Choosing the right place to start is always tricky, so rotating the protein allows you to see the full 3D shape you have existing, the colour of the segments are ideal when they are green, if the backbone colour is brown, this area can be rebuilt to find a new position that gives a better score “or” lower score initially, but the protein sits in a better position, by shaking and wiggling the protein will mean you can see how it’s movement and new rebuild holds together or just the area you rebuilt.

#### Modifying approx. three at a time will give you an indicator if you are working on the right spot

Sometimes having the amino acids in a different position and rebuilding those segments can help find new positions that weren’t available in the existing position. As the backbone is changed, it gives the possibility to find new energy levels that reflect the bonds to the surrounding amino acids within the existing structure. Some attract, some repulse and others are neutral to one another. Once the backbone has settled, you will be able to identify any areas that have changed from Brown to Green or Vice Versa, thus working on connected areas (segments) around the protein that “look” like they could be in a different combination can be tried.

Often when an amino acid or more are out of place, you can see that it’s a brownish colour and by manipulating the backbone into a new position will drop the energy level but once you have Shaked or Wiggled it will now take into consideration the new positions and by reducing the clashing importance lower you can make the backbone relax then adjusting the clashing importance higher it will give you some movement and it will explore the new environment you have given the protein to wiggle out.

With the Electron Density Reconstruction puzzles you will get some possibility areas that have a exploration area that are ghosted over top of the existing protein, you can try and rebuild the backbone to fit into this exploration area and after each adjustment, shake and wiggle out the protein to see how each little movement effects the score and shape of the protein.

If you try and make adjustments BEFORE wiggling the protein out, rebuild a few segments first, then decide whether to shake and or wiggle because the protein is usually hard to move much for these Puzzles.

Some of the amino acids will give you an idea of where in the protein structure they prefer to be, start or end of helix or the sheet, as you get familiar with different aminos and the backbone structure along with segment colours you will be able to choose a better configuration to hunt for better potential and find a shape that’s more natural.

### grogar7

I start by doing a whole-protein shake and wiggle at ci=0.2 to start. Then, with these larger proteins, I trim to the worst scoring areas and rebuild. Then I revert to the standard techniques I have described in other communications with you.

## References

1. H. Berman, K. Henrick, H. Nakamura, Announcing the worldwide Protein Data Bank. Nature Structural Biology 10, 980–980 (2003).

2. J. Jumper et al., Highly accurate protein structure prediction with AlphaFold. Nature 596, 583–589 (2021).

3. R. P. Joosten, T. Womack, G. Vriend, G. Bricogne, Re-refinement from deposited X-ray data can deliver improved models for most PDB entries. Acta Crystallogr D 65, 176–185 (2009).

4. R. J. Read et al., A new generation of crystallographic validation tools for the protein data bank. Structure 19, 1395–1412 (2011).

5. R. P. Joosten, G. Vriend, PDB Improvement Starts with Data Deposition. Science 317, 195–196 (2007).

6. B. van Beusekom et al., Homology-based hydrogen bond information improves crystallographic structures in the PDB. Protein Sci 27, 798–808 (2018).

7. R. P. Joosten, K. Joosten, G. N. Murshudov, A. Perrakis, PDB-REDO: constructive validation, more than just looking for errors. Acta Crystallogr D 68, 484–496 (2012).

8. S. Cooper et al., Predicting protein structures with a multiplayer online game. Nature 466, 756–760 (2010).

9. J. K. Leman et al., Macromolecular modeling and design in Rosetta: recent methods and frameworks. Nat Methods 17, 665–680 (2020).

10. S. Horowitz et al., Determining crystal structures through crowdsourcing and coursework. Nature Communications 7, (2016).

11. F. Khatib et al., Building de novo cryo-electron microscopy structures collaboratively with citizen scientists. Plos Biol 17, (2019).

12. V. B. Chen et al., MolProbity: all-atom structure validation for macromolecular crystallography. Acta Crystallogr D Biol Crystallogr 66, 12–21 (2010).

13. R. P. Joosten et al., A series of PDB related databases for everyday needs. Nucleic Acids Res 39, D411–419 (2011).

14. J. Cheng et al., Accurate proteome-wide missense variant effect prediction with AlphaMissense. Science 381, 1303-+ (2023).

15. D. Liebschner et al., Macromolecular structure determination using X-rays, neutrons and electrons: recent developments in. Acta Crystallogr D 75, 861–877 (2019).

16. P. V. Afonine et al., FEM: feature-enhanced map. Acta Crystallogr D 71, 646–666 (2015).

17. R. P. Joosten, F. Long, G. N. Murshudov, A. Perrakis, The PDB_REDO server for macromolecular structure model optimization. Iucrj 1, 213–220 (2014).

18. G. N. Murshudov et al., REFMAC5 for the refinement of macromolecular crystal structures. Acta Crystallogr D Biol Crystallogr 67, 355–367 (2011).

19. A. A. Vagin et al., REFMAC5 dictionary: organization of prior chemical knowledge and guidelines for its use. Acta Crystallogr D 60, 2184–2195 (2004).

